# Effect of Socio-environmental Factors on Rat Proliferation and *Leptospira* Transmission in Suburban Areas of Dungun, Terengganu, Malaysia

**DOI:** 10.1101/2023.07.13.548799

**Authors:** Mohammad Izuan Mohd Zamri, Nur Juliani Shafie, Mohammad Ridhuan Mohd Ali, Adedayo Michael Awoniyi, Hernan Dario Argibay, Federico Costa

**Author notes:** Correspondence to: NJS; FC.

## Abstract

**Background:** Rats play a significant role as important reservoirs in the transmission of neglected zoonotic diseases, such as leptospirosis, particularly in poor urban and suburban communities of Low and Middle-Income Countries. Their proliferation is influenced by poor socio-environmental conditions like poor housing conditions, improper refuse disposal, and open sewer which generally furnish rats with food, water and harborage sources. Several interventions have been targeted against rats, given their public health menace but with no significant success, probably due to insufficient knowledge about context-specific factors influencing their proliferation in poor environments.

**Methodology/Principal Findings:** We evaluated the effect of different socio-environmental conditions on rat proliferation and the possible outcome on the transmission of *Leptospira* in suburban environments. We performed ten trapping sessions between April 2021 and January 2022, and captured 89 small mammals from 1385 trapping efforts (specifically, *Rattus norvegicus* (n = 39), *Rattus rattus* (n = 27), *Rattus exulans* (n = 10), *Suncus murinus* (n = 11), and *Tupaia glis* (n = 2)), with a 15.7% (n = 14/89) of the captured animals testing positive for *Leptospira* bacteria using PCR detection. We used a generalized linear model to evaluate the effect of different socio-environmental variables on household rat infestation and reported residences without paved floors, the presence of animals/pets, residence type and residences with vacant lots as variables positively associated with rat proliferation in the study sites.

**Conclusion:** Rats are widely and differentially distributed in the poor communities of Dungun. Besides *R*. *norvegicus* and *R*. *rattus*, *S. murinus and T. glis* could also maintain and encourage pathogenic *Leptospira* transmission in the study areas. To adequately control rats proliferation and subsequent human zoonoses transmission, it will be critical to advocate and promote appropriate infrastructure and urban services, together with good hygiene practices to reduce rats’ access to water, food and harborage.

**Author Summary:** Rats are important reservoirs in the transmission of urban and suburban leptospirosis, and their proliferation is generally supported by poor socio-environmental conditions. Despite the strong association between poor socio-environmental conditions, rat proliferation and leptospirosis transmission, few studies have characterized their relationships in many poor suburban environments. Therefore, we evaluated the effect of different socio-environmental conditions on rat proliferation and possible effect on leptospirosis transmission in suburban communities of Dungun, Malaysia. We performed rat trapping and captured 89 small mammals from three orders, with a 15.7% (n = 14/89) of the captured animals testing positive for *Leptospira* bacteria using PCR detection. Using a generalized linear model, we reported residences without paved floors, the presence of animals/pets, residence type and residences with vacant lots as variables that are positively associated with rat proliferation in the study sites. Our findings show that rats are widely distributed in the study sites, and that in addition to *R*. *norvegicus* and *R*. *rattus*, *S. murinus and T. glis* could also maintain and transmit pathogenic *Leptospira*. Given the variables associated with rats proliferation, it is crucial to promote appropriate infrastructure and good hygiene practices to reduce rats’ access to water, food and harborage and subsequent rodent-borne diseases.

## Introduction

Leptospirosis, a bacterial disease that is caused by bacteria of the genus *Leptospira* affects approximately one million people worldwide annually, with most cases concentrated in Latin America and Asia, and an under-reported burden in Africa [1]. The bacterial disease is capable of infecting both humans and animals, and it is spread through the urine of infected animals, mostly rats [2]. According to Paul [3], *Leptospira* bacteria can be transmitted via direct or indirect routes and through droplets. Direct transmission occurs through contact between contaminated urine and human skin, i.e. abrasions, mucous membrane or other soft tissues [4], while indirect transmission occurs through contact between a contaminated surface such as water, soil and mud and human exposed parts like wounds, cuts and mucous membranes [5]. Infection may also occur via the consumption of contaminated food or water [6] or through the air, that is, when contaminated urine evaporates into the air and is inhaled by humans [7].

The propagation of rats, an important reservoir of *Leptospira* in urban and suburban environments, and the subsequent transmission of leptospirosis in most Low and Middle-Income Countries (LMICs) are attributable to a variety of socioeconomic and environmental conditions, among other factors [8]. For example, the majority of urban dwellers reside in regions that are characterized by low socioeconomic status, abandoned structural properties, poor household conditions, and improper refuse disposal, and as a result, provide rats with resources (i.e. water, food, and harborage sources) that facilitate their population proliferation and probable transmission of zoonoses in poor environments [9, 10].

Rats have been described as reservoirs for several zoonotic diseases, including bubonic plague, hantavirus, leptospirosis, toxoplasmosis, Lassa fever, and salmonellosis [11, 12]. In addition, rat-human cohabitation may affect human mental health through rat bites, disturbed sleep, allergies, irritation, social discrimination, and psychological trauma [13]. Besides the health implications, rat proliferation has been reported to be associated with significant agricultural loss, reduced economic productivity, ecosystem disruption, and household destruction, especially in poor LMICs urban environments [14, 15]. Given the current rate of globalization, there are growing concerns that the urban and suburban environments may become the hub of rat proliferation and associated risks since these areas promote conditions that favor rat population expansion [9].

Given the menace of rat proliferation, various control efforts or interventions are often implemented in affected areas to reduce their populations and in a way prevent possible disease transmission, and improve human public health [16]. Several rodent control methods have been used across the globe, but with mixed results [17]. For example, the chemical method, the easiest and most commonly used method, has demonstrated short-term effectiveness [18], while the combination of the chemical control method, sanitation improvement, and infrastructural intervention has somewhat shown a more promising result [19], but not the “golden control method”. Likewise, a combination of other methods like animal trapping and enhanced sanitation has also been proven to reduce rat infestations in poor environments [20]. However, none of these methods offers the desired long-term “golden control method”, as rats possess a high reproduction capacity, the ability to adapt to diverse habitat types, and the ability to easily migrate up to 90m from a neighboring colony even in poor urban environments amidst satisfactory food, water, and harborage sources [21], thereby repopulating treated areas after a period of successful intervention.

Perhaps, one of the reasons why effective and sustainable control of rats and indirect control of urban leptospirosis keep eluding us is in part due to our insufficient knowledge of the context-specific socio-environmental factors influencing rat population bloom in urban and suburban environments. This could be fairly responsible for the recent increase in the number of leptospirosis cases among the urban and suburban dwellers of Malaysia [22–24], where little is known about the population proliferation of rats in these environments and the specific social and environmental variables contributing to the spread of rats and the possible human-rat interactions and spread of zoonotic diseases, especially leptospirosis, in the poor LMICs urban environments. Previous studies conducted on *Leptospira* transmission in poor urban and suburban areas of Malaysia have primarily focused on the description of human leptospirosis [25], rodents and *Leptospira* spp. diversity [26, 27], and knowledge, attitude, and practice among the locals [28], with few or no studies adequately evaluating the relationship between the socio-environmental conditions influencing the proliferation of rats, the resultant possible high human-rat contact and pathogenic *Leptospira* transmission in poor urban and suburban environments of Malaysia. Therefore, this study aims to examine the socio-environmental factors contributing to rat proliferation and the probable transmission of *Leptospira* in poor environments characterized by increasing human leptospirosis cases. By so doing, it seeks to identify the key factors that may provide holistic solutions to the problem of rat infestation and consequent zoonoses transmission in poor urban and suburban environments.

## Methodology

### Study area

The study was conducted in the periphery of Dungun (4° 45’N 103° 25’E), Terengganu, Malaysia (Fig 1). Dungun is characterized by a typical monsoon season that normally occurs between November and December. Dungun is made up of 45% forest reserves, 35% lowland regions, 20% swamps, and other water bodies, with an estimated area of 2,735.03 km^2^ and a human population of about 193, 300 people [29]. Dungun was chosen for this study because of its high record of rodent sightings and cases of leptospirosis [30].

**Figure 1:**
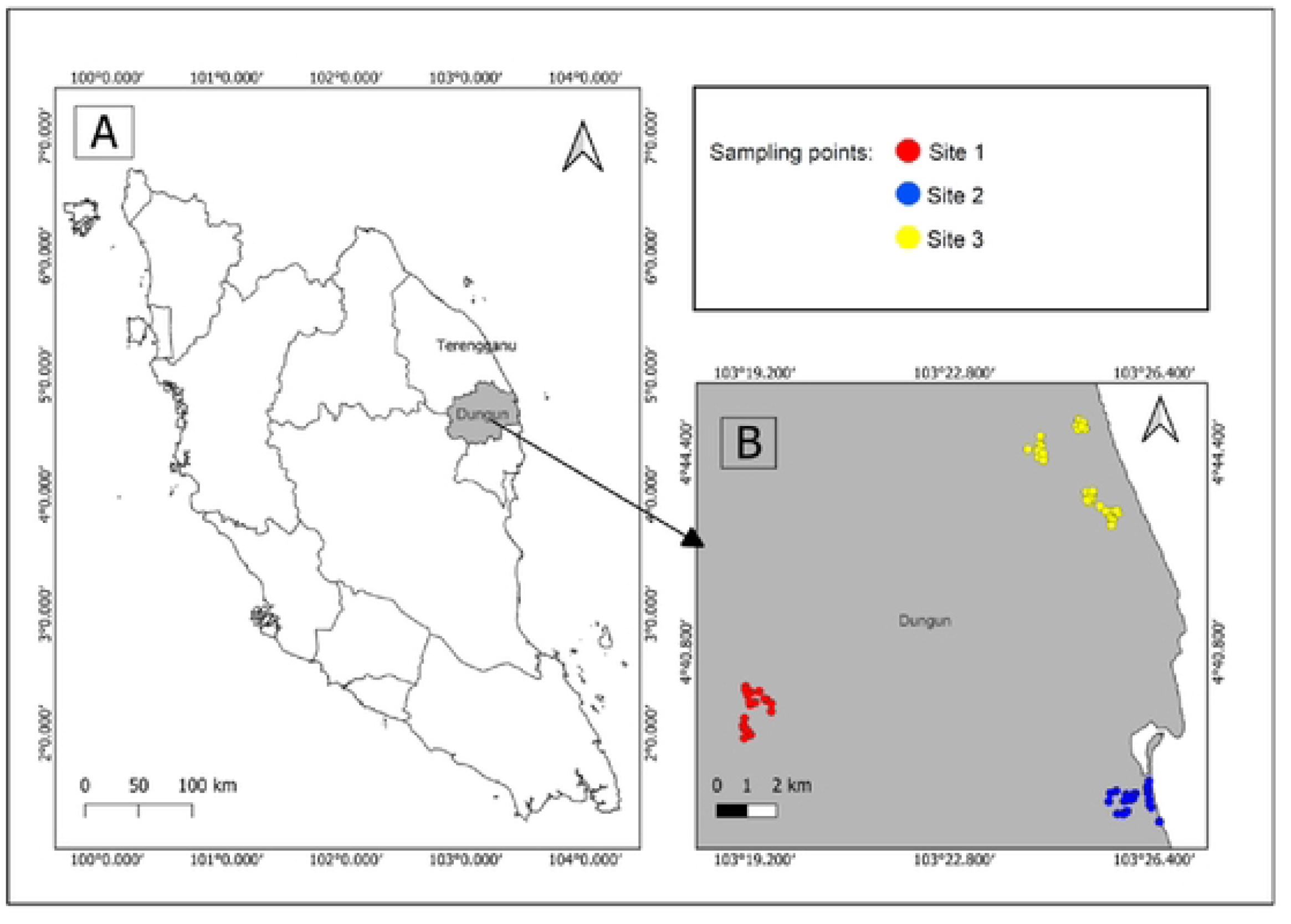
Map of Dungun, Terengganu in Malaysia showing the location of the study sites for rodent surveillance.

Briefly, the three study sites are located on the periphery of Dungun town. Specifically, site 1 is situated about 26 km from Dungun town. It constitutes scattered human settlements with some accompanying gardens in the backyards, livestock and oil palm plantations in the surrounding areas. The basic facilities on this site are road networks, schools, mosques, public offices, and several commercial buildings. Site 2 is located 20 km away from Dungun town and close to Paka Beach, a popular recreational centre in Dungun town. The site also possesses jetties for fishermen and wet markets where they trade their harvest. Other facilities on this site include residential areas, schools, and commercial buildings. The third site is located closer to Kuala Dungun town and possesses more basic facilities than the other two sites. This site has well-structured residential areas, medical facilities, supermarkets, and schools, but also a few poor scattered houses across the site. Also, site 3 has a low elevation with proximity to the river. This site is vulnerable to flooding, and usually experiences flooding after heavy rainfall [31].

### Study design

One hundred and nineteen sampling points were randomly selected from the three sampling sites (sites 1, 2 and 3) in a cross-sectional study over a period of ten months (April 2021 to January 2022).

### Evaluation

#### Animal trapping

We slightly modified the methods previously used and described by Awoniyi, Souza, et al. [2021] and Porter et al. [32] to trap rats using cage wire traps (25 cm x 15 cm x 12 cm). Briefly, we set up traps in and around houses, especially in areas with active rat signs, to increase our capture rate. On average, we set up at least 10 sausage-baited live traps at each point for three consecutive nights. To preserve the freshness of baits, traps were activated at dusk and checked the following morning, while baits of traps without animals were changed daily. All captured animals were maintained in traps and transported to the laboratory for further analysis. Captured animals were identified to the species level using the procedure previously validated by Francis and Barrett [33]. Kidneys of captured animals were harvested after an overdose of sodium pentobarbital anaesthesia, stored in a sterile specimen container and placed in a Panasonic Ultra-Low Freezer at −86℃ for further investigation. All animal operations were performed in accordance with the previously published protocol [34]. The Ethics Review Committee Board of the Universiti Malaysia Terengganu approved the method with approval number: UMT/JKEPHMK/2021/53.

#### Socio-environmental evaluation

Environmental surveys of the sampling points were conducted using a modified environmental survey form by Awoniyi et al. [19] and CDC [35]. The modified form includes the following six groups of variables: a) area type and ownership status, b) sources of food for rats, c) water sources, d) harborage sources, e) entry/access for rats, and f) active rat infestation signs, in addition to other relevant information. The questionnaires were administered to the head of household and standardized following the recommendations of Zahiruddin et al. [36].

### Pathogen investigation

#### DNA extraction

Following the manufacturer’s recommendations, we used the Macherey-Nagel NucleoSpin Tissue Genomic DNA Purification kit to extract at least 25 mg of kidney tissues from captured rats for initial DNA material. We performed modification on the final step and used 50 µl of pre-warmed TE buffer to dilute the DNA. The quantity and quality of the extracted DNA were examined using a Multiskan GO spectrophotometer. DNA from *Leptospira interrogans* serovar Pomona was diluted to 10 ng µL^-1^ to serve as a positive control in the subsequent Taqman qPCR, and genomic DNA was stored at −20 °C when required.

#### Taqman qPCR

PCR reactions were prepared to contain 1x GoTaq® Probe qPCR Master Mix, 400 nM of each forward and reverse primers, 200 nM of the probe, 200 nM of internal control primers, 150 nM of internal control probe, 8 µL of DNA template and PCR-grade water was adjusted to 20 µL [37]. For positive and NTC, 2 µL of leptospiral DNA (10 ng µL^-1^) or dH_2_O were used respectively. The PCR reactions were subjected to an amplification program which consisted of (i) an initial denaturation at 95 °C for 2 minutes, followed by (ii) 45 cycles of denaturation at 95 °C for 15 seconds and annealing and extension at 60 °C for 1 minute, using Biorad CFX96 thermal cycler. Any samples with sigmoidal curves passing the baseline threshold of 50, at Cq value ≤ 40 were considered positive.

### Statistical analysis

We used descriptive analysis to estimate trap success (precisely, we multiplied the total number of trapped rats by 100 and divided it by the total number of trapping efforts, while accounting for lost and damaged traps and those with non-targeted species) and the percentage of *Leptospira* positive rats. We used Chi-squared with Yates’s correction [38] to statistically test the difference between the number of captured, positive animals and the sites where the animals were captured, likewise the difference between the *Leptospira* positive animals and the species of captured animals. Also, we used generalized linear models (GLM) to examine the relationship between rat occurrence (present/absent) and the socio-environmental variables. Our response variable at each sampling point was coded as 1 if at least one active rat sign was observed or if at least one rat was captured during the three nights of rat trapping, or else coded as 0. Before testing for the relationship between rat occurrence and the socio-environmental variables in a multivariate GLM, in order to reduce the number of predictive variables to be included in the multi-factor model, we first used a separate GLM to test for the relationships between the response variable and each of the following explanatory variables: educational level of the head of household; type of the residence (residential/commercial); residence with backyard; residence bordering vacant lot; residence bordering open sewer; residence with unapproved trash storage; residence with fruits on the floor; water accessibility to rat; residence with unkempt bush; residence with accumulated waste; residence with dilapidated fence; residence with paved floor and holes on the walls; residence with pets; residence bordering an abandoned property. Variables with *p*-values of ≤ 0.2 from the single multi-factor model were included in a provisional multivariate model, since opting for the more conventional level of 0.05 at this stage could fail to identify all the important variables [39]. We then used a mixed forward and backward stepwise model selection approach to determine the final model using Akaike’s Information Criteria (AIC) [40]. The final model was the one with the lowest value of AIC, but the simplest model with a ΔAIC < 2 compared to this model was then selected based on the parsimony principle [41]. For all models, samples with missing values for any of the variables under consideration were excluded during the analysis. All analyses were performed in R version 4.3.0 [42].

## Results

### Rodent infestation

We found a noticeable rat infestation across the three study sites (Table 1), with a total of 89 small mammals captured during the 1385 total trapping efforts (6.43% trapping success). Five species, including three species of rodents (n = 76, 85.4%), one shrew (n = 11, 12.4%), and one treeshrew (n = 2, 2.2%) were captured. While we reported an apparent disparity in the diversity of small mammals captured across the study sites (Supplementary Material I), overall, *R. norvegicus* was the most abundant captured species (n = 39), with *T. glis* having the least number of captured species (n = 2) (Table 1). Also, the number of animals captured differed significantly across the study sites (*p* < 0.001), with site 2 having the highest number of captured individuals (n = 46), and site 3 recording the lowest number of captured animals (n = 20).

**Table 1.**
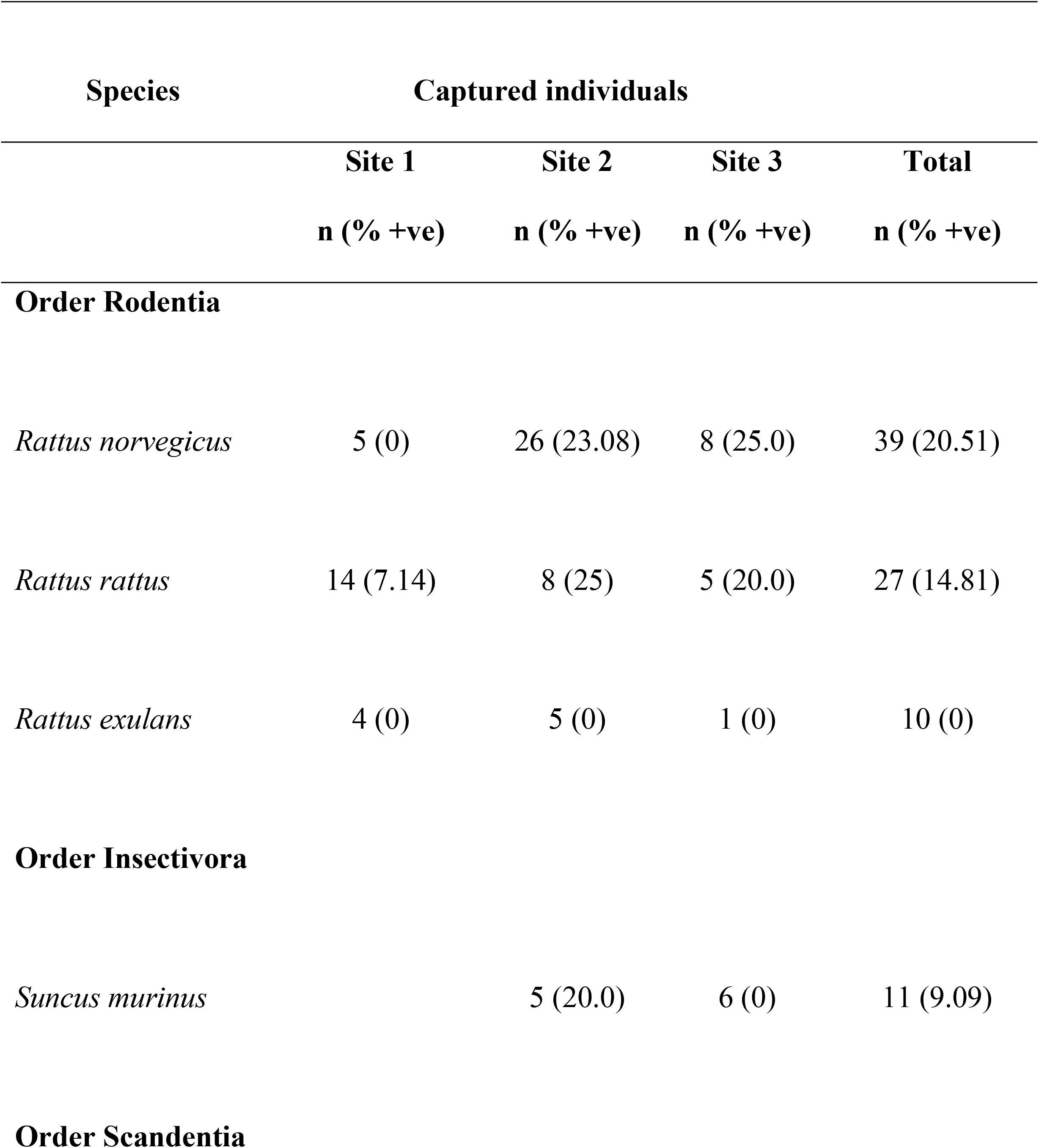
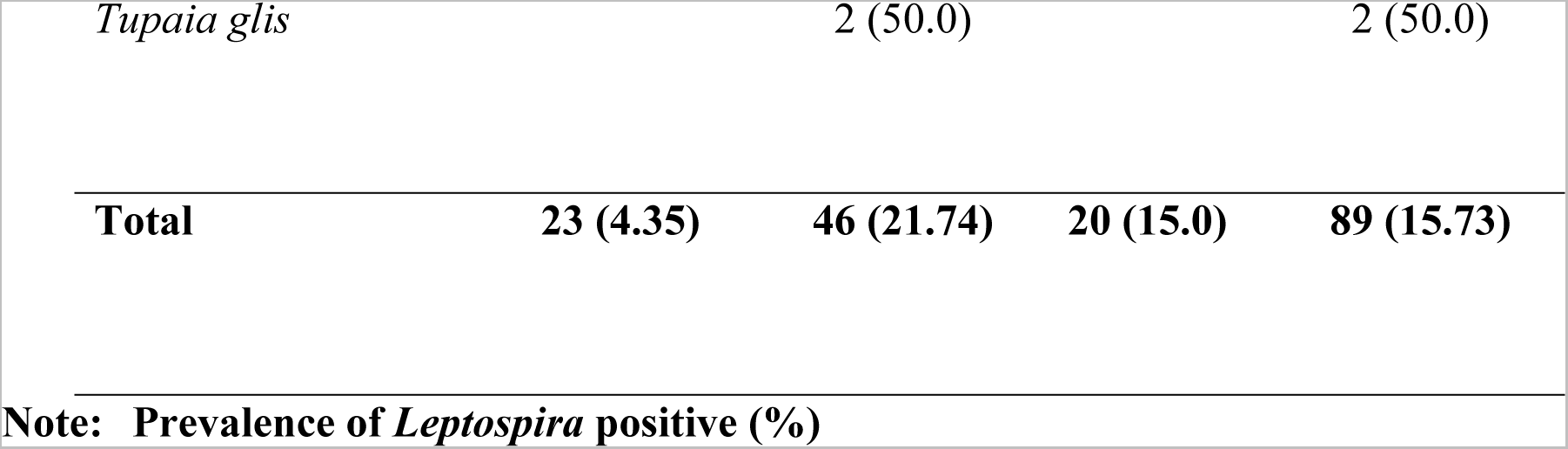
Distribution of captured small mammals and *Leptospira* positive individuals by study site and species.

#### *Leptospira* transmission

Out of 89 small mammals captured and screened for the pathogenic *Leptospira*, 14 (15.73%) tested positive for pathogenic *Leptospira*, amid significant variations across the study sites (*p =* 0.018). While all five species of small mammals but one tested positive for pathogenic *Leptospira* (Table 1), the result shows an apparent distribution of pathogenic *Leptospira* across the study sites, with the highest prevalence recorded in site 2 (21.74%), and *T. glis* (50%), and *R. norvegicus* (20.51%), the two species with the highest number of positive animals. Notably, out of the 10 individuals of *R. exulans* examined for pathogenic *Leptospira*, none tested positive across the three sites, while 1 of the 2 sparingly distributed *T. glis* (50%) tested positive for pathogenic *Leptospira*. However, Pearson’s Chi-squared test showed significant differences between the captured sites and PCR *Leptospira*-positive animals (χ² = 19.92, df = 9, *p* < 0.05).

### Factors associated with rodent proliferation

Fifteen variables (number of inhabitants of a household, education level of the head of household, residence type, residence with a backyard, residence bordering a vacant lot, residence with an abandoned vehicle, residence with unkempt surroundings, residence with accumulated waste, residence with construction materials, residence with a dilapidated fence, residence with a paved floor, residence with holes in the walls, abandoned residence, presence of animals and vegetation) with *p*-values of ≤ 0.2 from the initial analysis were considered for the generalized linear models. The final model retained five of these fifteen variables (residence with a paved floor, presence of animals, residence type, residence bordering a vacant lot, and presence of vegetation) with an AIC of 173.6, while the next best model had an AIC of 174.1 respectively. Residences with paved floors, the presence of animals, residence type, and residences bordering a vacant lot were positively associated with the proliferation of rodent infestations (Table 2 and Supplementary material II).

**Table 2.**
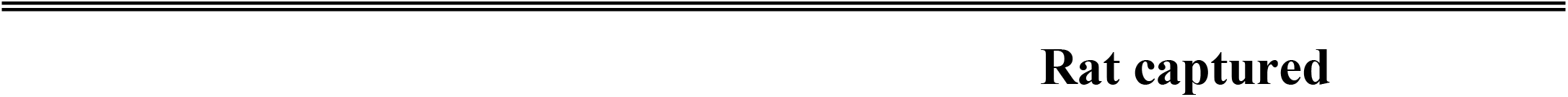
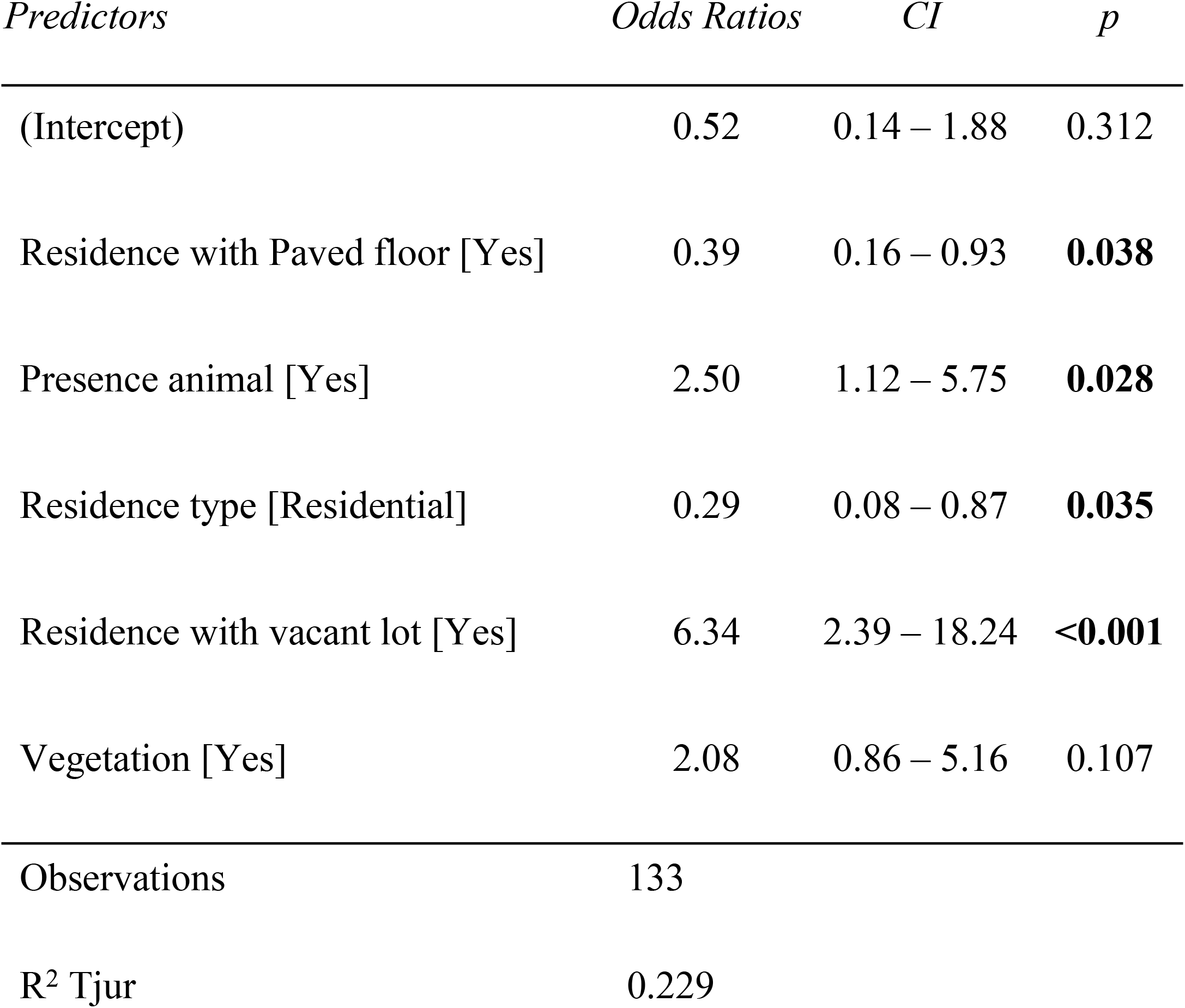
Summary of the generalized linear model showing variables that are associated with rodent infestation in the study sites.

**Rat captured**
| *Predictors* | *Odds Ratios* | *CI* | *p* |
| --- | --- | --- | --- |
| (Intercept) | 0.52 | 0.14 – 1.88 | 0.312 |
| Residence with Paved floor [Yes] | 0.39 | 0.16 – 0.93 | **0.038** |
| Presence animal [Yes] | 2.50 | 1.12 – 5.75 | **0.028** |
| Residence type [Residential] | 0.29 | 0.08 – 0.87 | **0.035** |
| Residence with vacant lot [Yes] | 6.34 | 2.39 – 18.24 | **<0.001** |
| Vegetation [Yes] | 2.08 | 0.86 – 5.16 | 0.107 |
| Observations | 133 |  |  |
| R<sup>2</sup> Tjur | 0.229 |  |  |

## Discussion

Our results show an apparent rat infestation and varied species diversity across the three study sites. The 6.43% trapping success recorded here is more than the 1.56% reported by Shafie et al. [27] from a recreational forest in Hutan Lipur Sekayu, Terengganu, Malaysia. The high trapping success recorded here could be a result of poor waste management regime, availability of food, water, and harborage sources across the study sites, as these have been shown to influence high rat proliferation [8], in addition to some residences bordering wet markets and oil palm plantations, which could provide rats with extra sources of food [24]. The significant disparity in captured species across the sites could be due to the varying environmental conditions noticed in the sites, for example, the settlements are disproportionately located near the beaches, wet markets, industrial warehouses, and mixed residential/commercial buildings, thereby differentially influencing the species distribution in the study sites with varying human-altered habitat. The higher number of species and captured animals recorded in study site 2 might indicate that this site possesses more impoverished conditions than the other study sites, for example, the wet markets could disproportionately attract rats with the fragrance of dry fish and other seafood and then provide them with food and harborage sources. Similarly, the higher abundance of *R. norvegicus* may be due to the species’ ability to swiftly adapt to almost all human-altered environments, with a tendency to dominate other rodent species [43].

Small mammals, especially rats, have been reported as the main reservoir of *Leptospira* in tropical and temperate environments [12], and their proliferation and high *Leptospira* prevalence (15.73%) in Terengganu show that rats are capable of maintaining and encouraging leptospirosis transmission in the study environments, particularly considering that almost all the *Rattus* species captured and tested in this study were positive for *Leptospira*. Although, we could only examine the presence of pathogenic *Leptospira* in two *T. glis,* the 50% prevalence of the bacteria in this species supports the report of Azhari et al. [44] that *T. glis* are also capable of influencing *Leptospira* transmission. Also, the long-distance travel capability of *T. glis* [45] can increase *Leptospira* transmission among other probable mammalian reservoirs in the environment, subsequently affecting public health by increasing the probability of human infections. The higher prevalence of pathogenic *Leptospira* in study site 2 could be associated with the higher distribution of rats recorded in this site. Although, the zero *Leptospira* prevalence recorded among *R. exulans* in this study is similar to the result of Blasdell et al. [46], who also recorded zero *Leptospira* prevalence among 3 *R. exulans*, further longitudinal investigation should be conducted to evaluate the potential of this species in the transmission of pathogenic *Leptospira*.

Based on our final model, in addition to residence floor status (that is, with or without paved floor), residences with the presence of animals, residence type (that is, residential or commercial), and residences with vacant lots (Supplemental Material II) had a significant association with rodent infestation in the study sites. These conditions are capable of providing rats with water, food, and harborage, which are necessary for their population sustenance and propagation in any environment [47]. These conditions are similar to those earlier reported to be positively associated with rodent infestation by Lambert et al. [48]. Also, residents residing close to restaurants, wet markets, and vacant lots reported high rat sightings during the survey, which might be due to the poor garbage disposal noticed at these points in addition to rats potentially obtaining harborage (overgrown vegetation, and debris) from the nearby oil palm plantation. This could encourage the proliferation of rats and other small mammals and subsequent frequent human-animal contact that may increase the probable circulation of pathogenic *Leptospira* in the study sites [24].

In conclusion, while we could not trap rats over a longer period (longitudinal study) to evaluate how seasonal variation potentially affects their population over time and the consequential effect on zoonotic prevalence, we have been able to provide useful information about the distribution of small mammals and their potential to transmit pathogenic *Leptospira* in suburban areas of Dungun, Terengganu, Malaysia. Our results illustrate that these animals, especially Norway rats and *T. glis* could maintain and encourage pathogenic *Leptospira* in the study areas. Given that few *T. glis* were captured and tested for the bacteria (with 50% of the tested animals positive for *Leptospira*), further studies should be conducted to evaluate the species reservoir potential to formulate a robust control plan. Likewise, more studies should also be conducted to characterize the role of *R. exulans* in the leptospirosis transmission chain. To adequately control rats proliferation and subsequent human zoonoses transmission, it will be critical to advocate and promote appropriate infrastructure and urban services, together with good hygiene practices to reduce rats’ access to water, food, and harborage.

## Acknowledgements

We want to thank the residents in Dungun, Terengganu for allowing us to conduct the animal trapping in their areas. We also want to extend our gratitude to the Ministry of Health Malaysia for granting the permission to conduct this study.

## Disclosure statement

No conflict of interest was declared by the authors.

## Funding

We thank Universiti Malaysia Terengganu for providing technical and funding support for this project (UMT/TAPE-RG/2020/55279).

## Authors’ contributions

Conceptualization: MIMZ; NJS; AMA & FC; Data curation: MIMZ; NJS; MRMA; AMA; HDA & FC; Formal analysis: NJS; AMA; HDA & FC; Funding acquisition: NJS; Methodology: NJS; MRMA; AMA & FC; Supervision: NJS; AMA & MRMA; Writing-original draft: MIMZ; NJS & AMA; and Writing-review & editing: NJS; AMA; HDA & FC.

## Data availability statement

All data and code used in this study is available in Zenodo under the Creative Common 4.0 license, accessible through https://doi.org/10.5281/zenodo.8023283.

## Notes

### Competing Interest Statement

The authors have declared no competing interest.

## References

1. Munoz-Zanzi C, Groene E, Morawski BM, Bonner K, Costa F, Bertherat E, et al. A systematic literature review of leptospirosis outbreaks worldwide, 1970-2012. Rev Panam Salud Publica. 2020; Jul 15;44:e78. doi: 10.26633/RPSP.2020.78

2. Levett PN. Leptospirosis. Clin Microbiol Rev. 2001; 14: 296–326

3. Paul NV. Clinical microbiology review. Leptospirosis. 2001; 14(2): 297

4. Bharti AR, Nally JE, Ricaldi JN, Matthias MA, Diaz MM, Lovett MA, et al. Leptospirosis: a zoonotic disease of global importance. The Lancet infectious diseases. 2003; 3(12):757–771

5. Ricaldi JN, Vinetz JM. Leptospirosis in the tropics and in travelers. Current infectious disease reports. 2006; 8:51–58.

6. Victoriano AFB, Smythe LD, Gloriani-Barzaga N, Cavinta LL, Kasai T, Limpakarnjanarat K, et al. Leptospirosis in the Asia Pacific region. BMC Infectious Diseases. 2009; 9:147. doi: 10.1186/1471-2334-9-147.

7. Heymann DL. Control of communicable diseases manual (No. Ed. 19). American Public Health Association. 2008

8. Costa F, Ribeiro GS, Felzemburgh RD, Santos N, Reis RB, Santos AC, et al. Influence of household rat infestation on leptospira transmission in the urban slum environment. PLoS Negl. Trop. Dis. 2014; 8(12): 3338. doi: 10.1371/journal.pntd.0003338

9. Felzemburgh RDM, Ribeiro GS, Costa F, Reis RB, Hagan JE, Melendez AXTO, et al. Prospective Study of Leptospirosis Transmission in an Urban Slum Community: Role of Poor Environment in Repeated Exposures to the Leptospira Agent. PLoS Neglected Tropical Diseases. 2014; 8(5). doi: 10.1371/journal.pntd.0002927

10. Leibler JH, Zakhour CM, Gadhoke P, Gaeta JM. Zoonotic and vector-borne infections among urban homeless and marginalized people in the United States and Europe, 1990–2014. Vector-Borne and Zoonotic Diseases. 2016; 16(7):435–444.

11. Battersby S. Rodents as carriers of diseases. In: Buckle A, Smith R, eds. Rodent pests and their control 2nd ed. Wallingford, Oxon UK: CABI International. 2015; 81–101. ISSN-13: 978-84593-817-8.

12. Costa F, Porter FH, Rodrigues G, Farias H, De Faria MT, Wunder EA, et al. Infections by *Leptospira interrogans*, seoul virus, and bartonella spp. among Norway rats (*Rattus norvegicus*) from the Urban slum environment in Brazil. Vector-Borne and Zoonotic Diseases. 2014;14(1): 33–40.

13. Byers KA, Cox SM, Lam R, Himsworth CG. “They’re always there”: resident experiences of living with rats in a disadvantaged urban neighbourhood. BMC Public Health. 2019; 19: 853 doi: 10.1186/s12889-019-7202-6

14. Belmain SR, Htwe NM, Kamal NQ, Singleton GR. Estimating rodent losses to stored rice as a means to assess ef fi cacy of rodent management. Wildlife Research. 2014; doi: org/10.1071/WR14189.

15. Pimentel D, Zuniga R, Morrison D. Update on the environmental and economic costs associated with alien-invasive species in the United States. Ecological Economics. 2005; 52(3): 273–288. doi: 10.1016/j.ecolecon.2004.10.002

16. Lambert MS, Quy RJ, Smith RH, Cowan DP. The effect of habitat management on home-range size and survival of rural Norway rat populations. Journal of Applied Ecology. 2008; doi: 10.1111/j.1365-2664.2008.01543.x

17. Oyedele DT, Sah SAM, Kairuddin L, Ibrahim WMMW. Range measurement and a habitat suitability map for the Norway rat in a highly developed urban environment. Trop. Life Sci. Res. 2015; 26(2): 27–44

18. Awoniyi AM, Thompson A, Ferguson L, Mckenzie M, Souza FN, Zeppelini CG, et al. Effect of chemical and sanitary intervention on rat sightings in urban communities of New Providence, the Bahamas. SN Applied Sciences. 2021; doi: org/10.1007/s42452-021-04459-x.

19. Awoniyi AM, Vargas CV, Souza FN, Zeppelini CG, Hacker KP, Pereira TC, et al. Population dynamics of synanthropic rodents after a chemical and infrastructural intervention in an urban low-income community. Scientific Reports. 2022; 1–10. Doi: 10.1038/s41598-022-14474-6.

20. Minter A, Costa F, Khalil H, Childs J, Diggle P, Ko AI, et al. Optimal Control of Rat-Borne Leptospirosis in an Urban Environment. Frontiers in Ecology and Evolution. 2019; 7: 1–10. doi: 10.3389/fevo.2019.00209

21. Awoniyi AM, Souza FN, Zeppelini CG, Xavier BI, Barreto AM, Santiago DC, et al. Using Rhodamine B to assess the movement of small mammals in an urban slum. Methods in Ecology and Evolution. 2021; 1–9. doi: 10.1111/2041-210X.13693.

22. Benacer D, Thong KL, Min NC, Verasahib KB, Galloway RL, Hartskeerl RA, et al. Epidemiology of Human Leptospirosis in Malaysia, 2004-2005. Acta Tropica. 2016; 157:162–168.

23. Benacer D, Thong KL, Verasahib KB, Galloway RL, Hartskeerl RA, Lewis JW, et al. Human Leptospirosis in Malaysia: Reviewing the Challenges after 8 Decades (1925-2012). Asia Pacific Journal of Public Health. 2016; 28: 290–302.

24. Garba B, Bahaman AR, Bejo SK, Zakaria Z, Mutalib AR, Bande F. Major epidemiological factors associated with leptospirosis in Malaysia. Acta tropica. 2018; 178:242–247.

25. Sahimin N, Sharif SA, Hanapi IRM, Chuan SN, Lewis JW, Douadi B, et al. Seroprevalence of anti-leptospira IgG and IgM antibodies and risk assessment of leptospirosis among urban poor communities in Kuala Lumpur, Malaysia. American Journal of Tropical Medicine and Hygiene. 2019; 101(6): 1265–1271. doi: 10.4269/ajtmh.19-0003

26. Pui CF, Bilung LM, Apun K, Su’ut L. Diversity of Leptospira spp. in rats and environment from urban areas of Sarawak, Malaysia. Journal of Tropical Medicine. 2017; 1–8.

27. Shafie NJ, Halim NSA, Awoniyi AM, Zalipah MN, Md-Nor S, Nazri MUIA, et al. Prevalence of Pathogenic Leptospira spp. in Non-Volant Small Mammals of Hutan Lipur Sekayu, Terengganu, Malaysia. Pathogens. 2022; 11: 1300. doi: 10.3390/pathogens11111300

28. Shafie NJ, Halim NSA, Zalipah MN, Amin NAZM, Esa SMAS, Md-Nor S, et al. Knowledge, Attitude, and Practices regarding Leptospirosis among Visitors to a Recreational Forest in Malaysia. The American journal of tropical medicine and hygiene. 2021; 104(4), 1290.

29. DMC - Official Portal of Dungun Municipal Council, Terengganu, Malaysia. 2022 https://mpd.terengganu.gov.my/index.php/ms/mpd/sumber/statistik (accessed 1 September 2022).

30. MOH-Ministry of Health Putrajaya, Malaysia. Distribution of leptospirosis cases in Terengganu, Malaysia. 2020; Unpublished data.

31. Ishak H, Sidek LM, Basri H, Fukami K, Hanapi MN, Lahat L, et al. Hydrologicalextreme flood event in Dungun River Basin region. In 13th International Conference on Urban Drainage. 2014; pp. 7–12 (PDF) FLOOD SIMULATION MODEL USING XP-SWMM ALONG TERENGGANU RIVER, MALAYSIA. Available from: https://www.researchgate.net/publication/319077066_FLOOD_SIMULATION_MODEL_USING_XP-SWMM_ALONG_TERENGGANU_RIVER_MALAYSIA [accessed Jul 11 2023].

32. Porter FH, Costa F, Rodrigues G, Farias H, Cunha M, Glass GE, et al. Morphometric and demographic differences between tropical and temperate Norway rats (*Rattus norvegicus*). Journal of Mammalogy. 2015; 96(2): 317–323. doi: 10.1093/jmammal/gyv033

33. Francis C, Barrett P. Guide to the mammals of Southeast Asia. 1st ed. United States: Princeton University Press. 2008.

34. Mills JN, Childs JE, Ksiazek TG, Peters CJ, Velleca WM. Methods for Trapping and Sampling Small Mammals for Virologic Testing. Services USDoHaH Ed. Atlanta, GA: U.S Dept. of Health & Human Service, Public Health Service, Centres for Disease Control and Prevention. 1995.

35. CDC. Integrated pest management: conducting urban rodent surveys. Centers for Disease Control and Prevention-Atlanta: US Department of Health and Human Services. 2006

36. Zahiruddin WM, Arifin WN, Mohd-Nazri S, Sukeri S, Zawaha I, Bakar RA, et al. Development and validation of a new knowledge, attitude, belief and practice questionnaire on leptospirosis in Malaysia. BMC Public Health. 2018; 18(1):331. doi: 10.1186/s12889-018-5234-y

37. Podgoršek D, Ružić-Sabljić E, Logar M, Pavlović A, Remec T, Baklan Z, et al. Evaluation of real-time PCR targeting the lipL32 gene for diagnosis of *Leptospira* infection. BMC Microbiology. 2020; 20(1):1–9.

38. Zar J. Biostatistical Analysis. Englewood Cliffs: Prentice Hall. 1996

39. Bursac Z, Gauss CH, Williams DK, Hosmer DW. Purposeful selection of variables in logistic regression. Source Code Biol. Med. 2008; 8:1–8 doi: 10.1186/1751-0473-3-17.

40. Akaike H. Factor analysis and AIC. Psychometrika. 1987; 52: 317–332.

41. Burnham KP, Anderson DR. Model selection and multimodel inference: A practical information-theoretic approach (Springer, 2002). 2002

42. R Core Team. R: A language and environment for statistical computing. R Foundation for Statistical Computing, Vienna, Austria. 2019; https://www.R-project.org.

43. Harper GA, Bunbury N, Invasive rats on tropical islands: their population biology and impacts on native species. Global Ecology and Conservation. 2015; 3: 607–627.

44. Azhari NN, Ramli SNA, Joseph N, Philip N, Mustapha NF, Ishak SN, et al. Molecular characterization of pathogenic Leptospira sp. in small mammals captured from the human leptospirosis suspected areas of Selangor state, Malaysia. Acta tropica. 2018; 188: 68–77.

45. Mariana A, Shukor MN, Muhd NH, Intan NB, Ho TM. Movements and home range of a common species of tree–shrew, Tupaia glis, surrounding houses of otoacariasis cases in Kuantan, Pahang, Malaysia. Asian Pacific Journal of Tropical Medicine. 2010;3(6): 427–434.

46. Blasdell KR, Morand S, Perera D, Firth C. Association of rodent-borne *Leptospira* spp. with urban environments in Malaysian Borneo. PLoS Negl Trop Dis. 2019; 13(2): e0007141. Doi: 10.1371/journal.pntd.0007141

47. Brooks JE. Methods of sewer rat control. In Proceedings of the 1st Vertebrate Pest Conference. 1962; https://digitalcommons.unl.edu/vpcone/17. Accessed 29 May 2023.

48. Lambert M, Vial F, Pietravalle S, Cowan D. Results of a 15-year systematic survey of commensal rodents in English dwellings. Scientific Reports. 2017; 7(1): 15882

